# Contrasting Response of Microeukaryotic and Bacterial Communities to the Interplay of Seasonality and Stochastic Events in Shallow Soda Lakes

**DOI:** 10.1101/2023.03.15.532723

**Authors:** Zsuzsanna Márton, Bianka Csitári, Tamás Felföldi, Ferenc Jordán, András Hidas, Attila Szabó, Anna J. Székely

## Abstract

Seasonal environmental variation is a leading driver of microbial planktonic community assembly and interactions. Yet, unexpected departures from general seasonal successional trends are often reported. To understand the role of local stochastic events in modifying seasonal succession, we sampled fortnightly throughout three seasons (spring, summer, and autumn) five nearby shallow soda lakes exposed to the same seasonal meteorological changes. We characterised their microeukaryotic and bacterial communities by 18S and 16S rRNA gene sequencing, respectively. Biological interactions were inferred by the analyses of synchronous and time-shifted interaction networks, and the keystone taxa were topologically identified. The pans showed similar succession patterns during the study period with spring being characterised by high relevance of trophic interactions and certain level of community stability followed by a more dynamic and variable summer-autumn period both in respect of community composition and microbial interactions. Adaptation to general seasonal changes happened through the abundant shared core microbiome of the pans. However, stochastic events such as desiccation and cyanobacterial blooms disrupted common network attributes and introduced shifts from the prevalent seasonal trajectory. These were more pronounced for microeukaryotes than for bacteria which was reflected in increased turnover and contribution of non-core microeukaryotes. Our results demonstrated that despite being extreme and highly variable habitats, shallow soda lakes exhibit certain similarities in the seasonality of their planktonic communities, yet random stochastic events such as droughts can instigate substantial deviations from prevalent trends for the microeukaryotic but not bacterial communities.

## Introduction

Seasonal changes of environmental parameters are the primary drivers of annual succession dynamics of planktonic communities instigating characteristic ecological processes repeated each year ^1,2^ and making seasonality the focal point of numerous aquatic microbial ecology studies ^3–5^. Departures from usual seasonal patterns are usually explained by interannual climatic variations or long-term trends ^6^. Seasonal changes affect shallow lakes in particular as they have high surface-to-volume ratio ^7–9^. Endorheic shallow lakes are even more impacted by external environmental changes due to their intrinsic lack of outflow and consequent importance of evaporation ^10–12^ making them ideal systems for the study of the seasonality of pelagic communities. Although the influence of interannual variation on the planktonic communities of endorheic aquatic systems has been shown before ^13,14^, it is not known whether such variations arise as a consequence of the interannual variations of the seasonal environmental dynamics or are driven by local stochastic events.

Soda lakes are a form of saline endorheic lakes characterised by carbonate, bicarbonate, and sodium as dominant ions ^15^. These systems represent the most alkaline aquatic habitats on the surface of our planet ^16–18^. The Kiskunság National Park (Hungary), in the central part of the Carpathian Basin (Central Europe) is a region with particularly high density of soda pans (i.e., shallow soda lakes) ^19^. The climate of this region is temperate continental ^20^ which creates considerable seasonal temperature and water level fluctuations. The drought prolonging and precipitation pattern modifying effect of climate change ^21–23^ is also substantial resulting, among others, in increasing frequency of desiccation events ^24^. It has been implicated that climate change induced weather anomalies also exacerbate the interannual variation in the seasonality of the biogeochemistry of individual pans ^10^. Nevertheless, while the seasonal environmental changes are becoming less predictable, the close proximity of these pans means that they are still exposed to practically identical climatic conditions. This makes the soda pans of Kiskunság a perfect system to disentangle the impact of local stochastic events on planktonic communities from the effect of the common seasonal environmental changes.

For aquatic microorganisms diversity alone is a usually poor predictor of functional capacities ^25,26^, which has been explained by the limitation of species composition-based diversity assessments in capturing the complexity of microhabitats and the associated microbial interactions ^27^. Microbial interactions are critical to both microbial functioning ^28^ and community assembly ^29^. Therefore, to understand the seasonality of aquatic microbial communities, it is important to focus not only on compositional changes but also on species interactions ^30^. Due to the complexity of microbial communities, network perspective is required to study their species interactions ^29^. Synchronous networks can assess the simultaneous occurrence and abundance of organisms, while time-shifted association networks can elucidate time-delayed impacts of species interactions ^6,31,32^. Network analysis can also identify keystone species ^6,32,33^ which are usually defined as hubs (i.e., highly associated species in the network) that have disproportionately high importance in the community relative to their abundance ^33–35^.

Although microbial communities typically consist of hundreds or thousands of species (Bengtsson-Palme, 2020), only a smaller fraction of the microbial species are shared among communities of a specific habitat type (e.g., soda lakes) and can be defined as its core microbiome ^36^. The core microbiome is hypothesised to represent the functionally or ecologically most important taxa of a habitat type ^37,38^. It has been also suggested that microbial adaptation to environmental changes through dispersal mediated species sorting also operates primarily on the core microbiome instead of the rare or sporadic (i.e., non-core) community members ^39^.

Soda lakes and pans are characterised by abundant phytoplankton ^13,40^ zooplankton ^41,42^ that feeds large amounts of migratory birds ^43,44^, while fish are typically absent ^45^. Studies describing the planktonic microbiome of soda pans in the Carpathian Basin ^45–48^, Canada ^49^, Brazilia ^50^, China ^51^ and East Africa ^52^ have identified unique bacterial and eukaryotic communities. The seasonal changes of the composition ^46,47^ and function ^53^ of bacterial communities of soda lakes and pans have also been described before, but the impact of seasonality on the interactions of bacterial and microeukaryotic species remains to be explored. Due to inherent biological differences such as cell size and structure, generation times, and life history traits, the bacterial and microeukaryotic communities are expected to respond differently to environmental changes including seasonality ^54–57^.

We carried out an extensive field study evaluating the seasonal dynamics of planktonic microbial communities of five soda pans located in the same region (i.e., Kiskunság NP) throughout three seasons (spring, summer, autumn) by fortnightly sampling. We aimed to understand the impact of seasonality on the structure and interactions of bacterial and microeukaryotic communities and the contribution of core and non-core microbiome to the adaptation to environmental changes. Our primary hypothesis was that common habitat type (i.e., shallow soda lake) and identical climatic exposure due to the proximity and synchronous sampling of the five sites results in analogous seasonal succession patterns both in respect of community composition and interactions. We further hypothesise that adaptation to seasonal changes happens to a larger extent through species recruitment from the core than the non-core community, while non-core taxa are primarily involved in the response to local stochastic events. Finally, we hypothesise that bacterial communities are less affected by stochastic events than eukaryotic communities.

## Materials and methods

### Sample collection and environmental parameters

We selected five soda pans located within an area of 14 km^2^ of the Kiskunság National Park, Hungary: Böddi-szék, Kelemen-szék, Sós-ér, Zab-szék and an unnamed pan (Pan no. 60) (Figure 1). Two different subtypes of soda pan are distinguished in this region: the prevailing turbid-white subtype (referred as turbid from here on), which is characterised by high amounts of suspended inorganic clay particles and the less common non-turbid, brown subtype (referred here as brown), which has characteristic brown colour due to high organic material content and low amount of suspended inorganic material ^19^. Among the sampled pans Sós-ér was brown, while the other pans were turbid.

The pans were sampled fortnightly from April 12 to November 14 in 2017, covering three seasons: spring (sampling time 1-4), summer (sampling time 5-10) and autumn (sampling time 11-14) (Figure 2a). Each time, composite samples were collected by sampling water from 1 cm below the surface at least at five different points near the deepest part of the pans. For microbial community analyses, water samples were collected into a 1-litre sterile bottle after filtering through a 40 µm mesh size plankton net to remove large zooplankton; larger algae are practically absent from these pans; the mean contribution of pico-sized (cell diameter <2 µm) algae to total phytoplankton biomass is ∼85% in the turbid pans ^40^, while in the brown soda pans nanoplanktonic (2-20 µm) algae have similar contribution (∼80%) with moderate amount of picophytoplankton (∼8%) ^40,47^. The water samples were transferred to the lab in a cooling box where upon arrival 30 ml water from the turbid soda pans and 50 ml from the brown pan were filtered onto 0.1 µm pore size filters (MF-Millipore membrane filter) to collect planktonic microbial cells. The filters were stored at – 80 °C until further processing.

**Figure 2.**
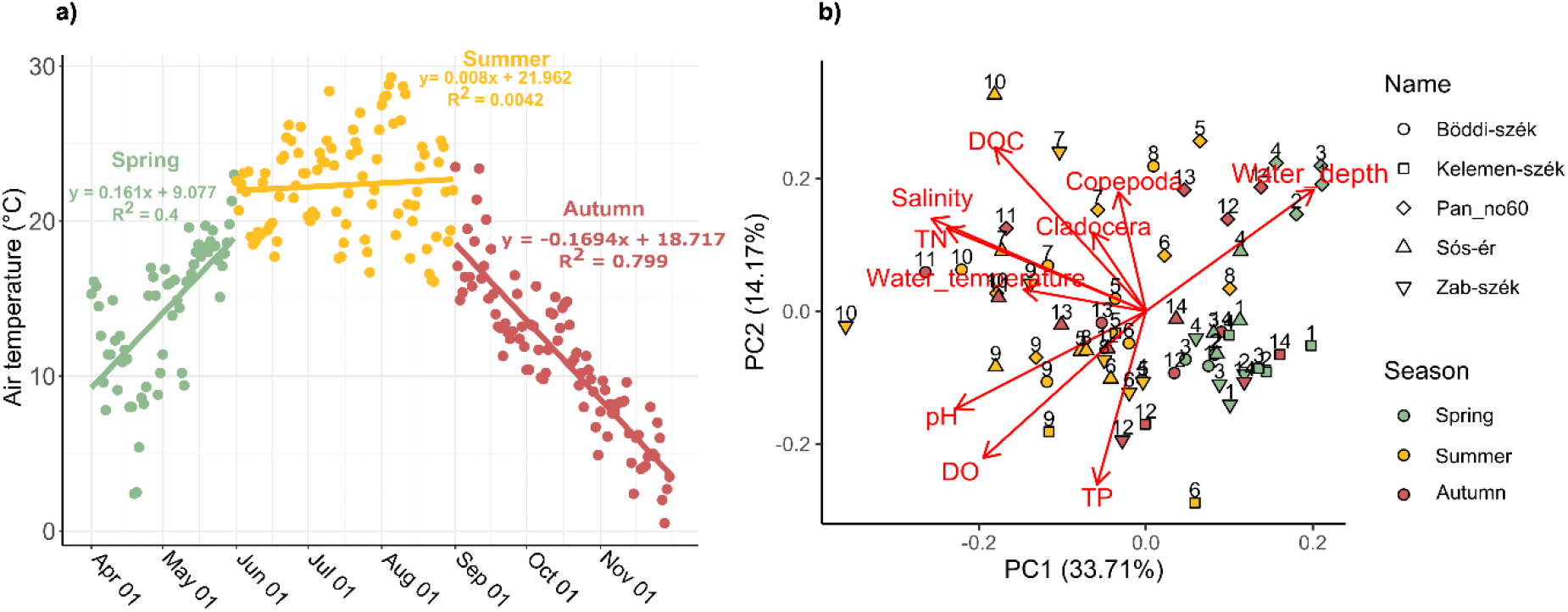
Seasonal changes. a) Air temperature trend in the sampling period, b) PCA biplot of the environmental variables of soda pan samples

Temperature, pH (SenTix 41 electrode), conductivity (TetraCon 325 cell) and dissolved O_2_ (CellOx-325 electrode) were measured on site with a MultiLine Handheld Meter model 340i (WTW, Weilheim in Oberbayern, Germany). Meteorological data from the Soltszentimre station (within 15 km distance from all sampling sites) was provided by the Hungarian Meteorological Service. Further environmental parameters were determined in the laboratory. Chlorophyll a (ChlA) was measured based on a previously published study ^58^, while total phosphorus (TP), total nitrogen (TN) and dissolved organic carbon concentrations were determined as described in Nydahl et al., 2019 ^59^. Conductivity data were converted to salinity based on a previously published equation ^19^. A total of 20 litres of water collected at different points around the deepest part of the open water area were sieved through a 40 µm mesh size plankton net and preserved in 70% ethanol until determination of abundance and composition of zooplankton ^60^.

### Community analysis

DNA was extracted using DNeasy PowerSoil Kit (QIAGEN) according to the manufacturer’s instructions. Microeukaryotic and bacterial community composition was determined by amplification and sequencing of the V4-V5 region of the 18S rRNA gene and V3-V4 region of the 16S rRNA gene, respectively (no Archaea specific PCRs were carried out, since they represent only a minor fraction of the prokaryotic community in these sites; ^47,61^). Amplification and library preparation for amplicon sequencing was carried out based on previously published studies^57,62^; https://www.protocols.io/view/sample-preparation-for-illumina-miseq-dual-index-a-badeia3e?step=1) (Text S1.), while sequencing was performed at the Swedish National Genomics Infrastructure (NGI) (Uppsala, Sweden) on Illumina MiSeq platform (Illumina Inc, San Diego, CA, USA) in a 2×300 bp paired-end format and with v3 chemistry.

Sequence read processing, alignment and taxonomic assignments were carried out using mothur v1.41.1 (Schloss et al., 2009). OTUs were assigned at 99% similarity cutoff and rarefied OTU sets were created as a basis of subsequent analyses. For the amplified region and based on previous studies of the surveyed habitat, this cutoff value was found suitable to avoid most diversity estimation biases caused by clustering different species to the same OTU or splitting organisms having multiple rRNA copies to separate clusters ^63,64^. The 7th sampling time of Pan no. 60 was discarded from the 18S rRNA gene amplicon dataset due to a low amount of high-quality sequences. Detailed description of sequence analysis is provided in supplementary Text S2.

OTUs present in all of the five studied soda pans were defined as core5 and OTUs shared between the four turbid soda pans as core4. OTUs not shared between the pans were defined as non-core5 and non-core4, respectively.

### Statistical analyses

Statistical analyses were performed using the “vegan” R package ^65^. The environmental variables were compared using principal component analysis (PCA). As a proxy of community turnover to assess structural temporal dynamics, Bray-Curtis dissimilarity between subsequent sampling occasions of each pan was calculated for both 18S rRNA and 16S rRNA gene OTUs (i.e., eOTUs and bOTUs, respectively). To compare the communities of the turbid and brown pans one-way permutational multivariate analysis of variance (PERMANOVA, “adonis” function, permutations = 999) based on Bray-Curtis dissimilarity of the eOTUs and bOTUS was performed, while differences in community compositions among seasons and lake identity were tested by two-way PERMANOVA (permutations = 999). Non-metric multidimensional scaling (NMDS) based on Bray-Curtis distance was used to visualise the microeukaryotic and bacterial communities, while “envfit” function was applied to plot significantly fitted (p < 0.05) environmental vectors onto the NMDS ordinations. One-way analysis of variance (ANOVA) with Tukey’s post-hoc test was used to test the differences of environmental variables between soda pans.

### Network analyses

The extended Local Similarity Analysis (eLSA) is a robust time series analysis tool that not only detects microbial associations that are present throughout the study period (i.e., global correlations) but also those that only occur in a subinterval of the time series (i.e., local associations). Furthermore, eLSA considers both associations between taxa that coexists in time (i.e., co-occurence) and time-shifted or time-lagged correlations ^6,66,67^. To better understand synchronous and asynchronous interactions, we generated two eLSA networks of the microeukaryotic and bacterial communities of each soda pan: one using the synchronous correlations (i.e., co-occurrence networks, delay 0) and another using only time-shifted correlations (delay 1 or -1). The eLSA was carried out for each pan using the default settings except for adjusting delay limit to 1 and data normalisation with the percentileZ function. To reduce the complexity, only OTUs with > 1% relative abundance in at least one sample and present with more than 10 reads in at least three different subsampled samples were included in the network analysis. For both global and local associations only strongly significant (p <0.01 and q <0.01) correlations were included (global: Spearman’s rank correlation coefficients (SSCC) and local similarity scores (LS)). Network visualisation was carried out with Cytoscape v3.8.2 using the edge-weighted spring-embedded layout.

In order to identify OTUs in important network positions and to quantify their centrality, we used the weighted topological importance (WI) measure suggested by Müller et al. (1999) and generalised by Jordán et al. (2006). This index calculates the number of neighbours and also the number of their neighbours, in an additive and multiplicative way while considering the strength of the interactions. We considered indirect interactions up to three steps (WI^3^). In order to differentiate the keystone OTUs (i.e., the OTUs that play a key role in the network and their removal would drastically impact the structure of the network) (Jordán et al., 2006; Berry & Widder, 2014), OTUs with WI_i3_ > 1 were selected. If less than six OTUs fulfilled this requirement in a network, the selection was expanded to include WI_i3_ ≥ 1 OTUs. In the network of interactions, one can speak of “negative keystone OTUs” that are richly connected via negative associations and, contrariwise, “positive keystone OTUs” that are rich in positive associations with others. We note that positive associations can be strongly transitive (AB and BC frequently implies AC), while negative associations are rarely transitive (AB and BC generates a positive AC association instead of AC in the negative network). Two heatmaps were generated using ‘‘ComplexHeatmap’’ R package ^68^ to visualise the z-score transformed abundance of the keystone OTUs.

## Results

### Environmental parameters

Weather data revealed typical seasonal air temperature dynamics with an increasing trend in spring (rate: +0.16 °C/day, mean: 14.1 °C), no trend in summer (mean: 22.3 °C) and decreasing trend in autumn (rate: -0.17 °C/day, mean: 11.3 °C) (Figure 2a). The measured environmental parameters had values and followed trends previously described for the soda pans of this region (Table S1) ^10,19,45,47^. In general, spring samplings had similar environmental conditions, while in summer and autumn variation increased both between sampling times and sites (Figure 2b, Figure S1). Water depth varied greatly (1.5-46.0 cm) during the sampling period with the deepest water levels measured during spring. Moreover Zab-szék and Kelemen-szék were completely dry at some occasions (sampling time 7, 8, 10, 11 and 13 for Kelemen-szék; and 11 and 13 for Zab-szék) making water sampling impossible. According to the PCA biplot, water depth was negatively related to the concentration of soluble compounds, pH, as well as copepoda and cladocera, which had the highest values in mid summer, while pH, DO and TP was elevated in summer and early autumn (Figure 2b, Figure S1). The measured environmental parameters clearly distinguished the brown Sós-ér, which had in average significantly deeper waters and higher TN concentrations than the turbid pans. For detailed description of the trends in environmental parameters, check Text S3.

### Community composition

A total of 4524 microeueukaryotic OTUs (eOTUs) and 4241 bacterial OTUs (bOTUs) were identified from the sequencing data. In all five lakes the three most abundant microeukaryotic phyla were Chlorophyta (mean relative abundance: 52.5%; range: 8.1-97.6%), Ochrophyta (14.9%; 0-82.4%) and Fungi (3.9%; 0-59.0%). Within the Chlorophyta phylum the single most abundant eOTU in all pans was a green algae affiliated to the *Choricystis* genus (abbreviated name Ch, 20.9%; 0.1-86.4%). This *Choricystis* eOTU showed clear seasonality with a mean abundance in spring of 45.7% which decreased to 9.4% in summer and only 3.0% in autumn, although by November it increased again to 6.9% (Figure 3a). Actinobacteria (30.4%; 7.0-69.3%) and Cyanobacteria (10.0%; 0-46.9%) were the most abundant bacterial phyla in all pans. Within Actinobacteria, the most abundant bOTU (Ni, 3.8%; 0-29.5%) belonged to the Nitriliruptoraceae family. Meanwhile OTUs identified as Cyanobium_PCC-6307 (Cy) and *Synechococcus*_MBIC10613 (Sy) were the most frequent cyanobacterial lineages (Figure 3b). The seasonal community dynamics of the brown Sós-ér differed from those of the turbid pans. In spring a bOTU belonging to the family Erysipelotrichaceae (Er) was dominant, while in the second half of summer a filamentous nitrogen-fixing *Nodularia*_PCC-9350 (No) cyanobacteria bOTU bloomed. Some eOTUs such as the members of Chrysophyceae Clades D (Cd) and F (Cf) or the parasitic fungus genus *Pythium* (Py) were only dominant (>1%) in Sós-ér (Figure 3). Meanwhile the turbid pans had relatively similar community dynamics, the only exception were the drastic shifts in the microeukaryotic community composition observed following the desiccation-refillment events in Kelemen-szék and Zab-szék when specific eOTUs became dominant (e.g. after the first desiccation in Kelemen-szék the ciliate *Halteria* (Hi) had 59.3% abundance, while in Zab-szék a Stramenopiles (St) eOTU became the most abundant).

**Figure 3.**
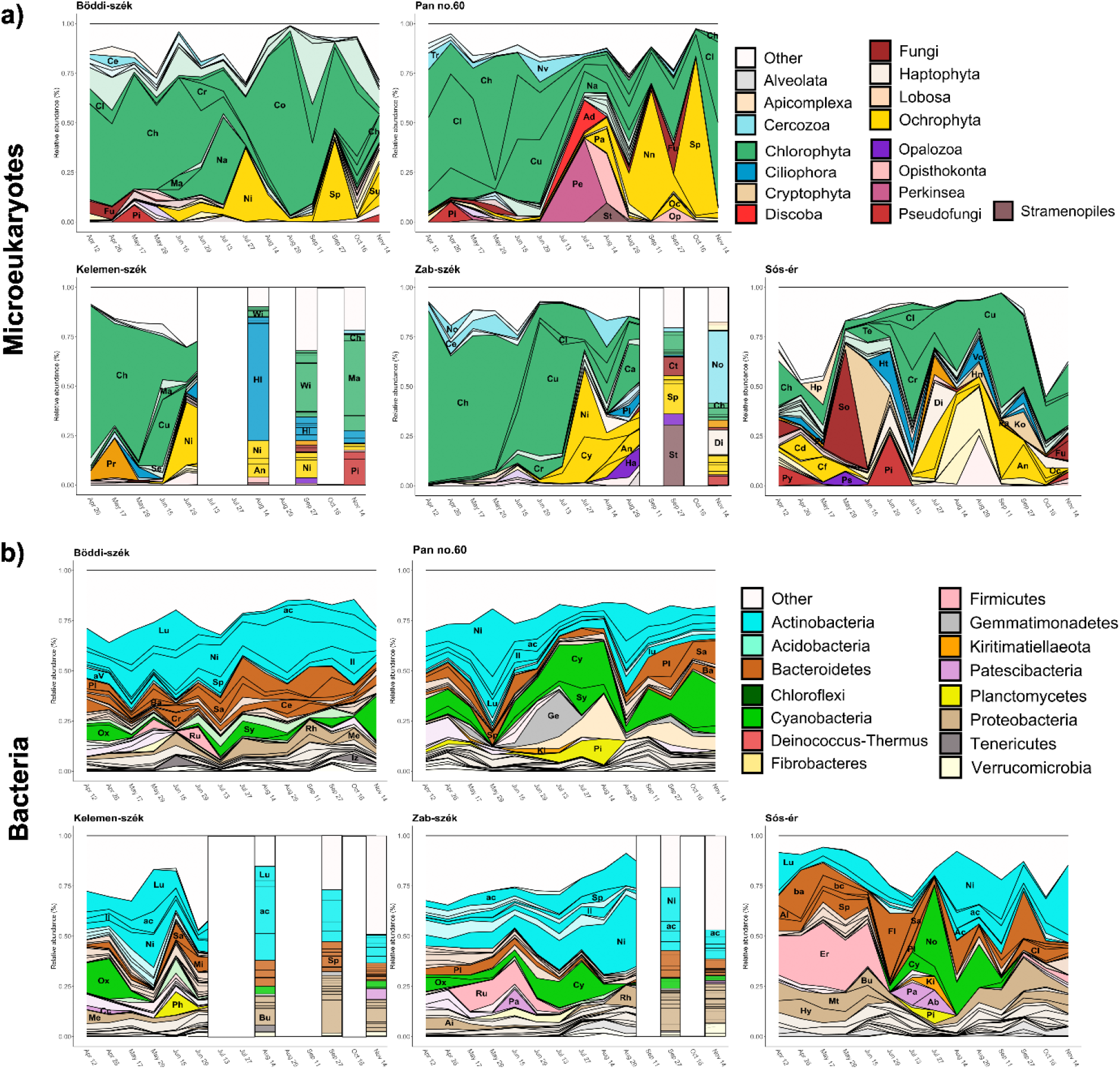
Microbial community dynamics of the a) microeukaryotic and b) bacterial OTUs with > 1% relative abundance in at least one sample of the given lake coloured according to the corresponding phyla. OTUs with > 5% relative abundance in at least one sample are highlighted with vivid colours and indicated by the abbreviated name of the corresponding microeukaryotic genera or bacterial clades, respectively. Key to the abbreviations can be found in the supplementary material Table S2.

### Drivers of community changes

The ‘envfit’ analysis significantly (p < 0.05) fitted salinity pH, DOC, TN, TP, and DO on the NMDS plots of both the microeukaryotic and bacterial communities. However, water depth and *Daphnia magna* abundance were significantly fitted only on the microeukaryotic NMDS, while water temperature, and chlorophyll were only significant for bacterial communities (Figure 4). The season of sampling had a significant effect on the communities (PERMANOVA Microeukaryotes: R^2^ = 0.144, p = 0.001; Bacteria: R^2^ = 0.115, p = 0.001) and the differences between seasons were also significant with the strongest differentiation of spring communities from summer and autumn (Table S3). Interestingly, the last autumn samples (sampling 14) appear close to the spring samples on both microeukaryotic and bacterial NMDS plots suggesting similarity between the spring and late autumn samples (Figure 4). According to the NMDS plots and the PERMANOVA analysis testing the impact of soda lake subtype (i.e., brown or turbid) both microeukaryotic and bacterial communities of the brown Sós-ér clearly separated from the communities of the other four turbid pans (Microeukaryotes: R^2^ = 0.086, p = 0.001; Bacteria: R^2^ = 0.145, p = 0.001). Not surprisingly, in the two-way PERMANOVAs including all five lakes and assessing the effect of lake identity and sampling season, lake identity always explained more variance than the season of sampling irrespective of the analysed domain (microeukaryotes or bacteria) and the subset of the community (all OTUs, core or non-core).

**Figure 4.**
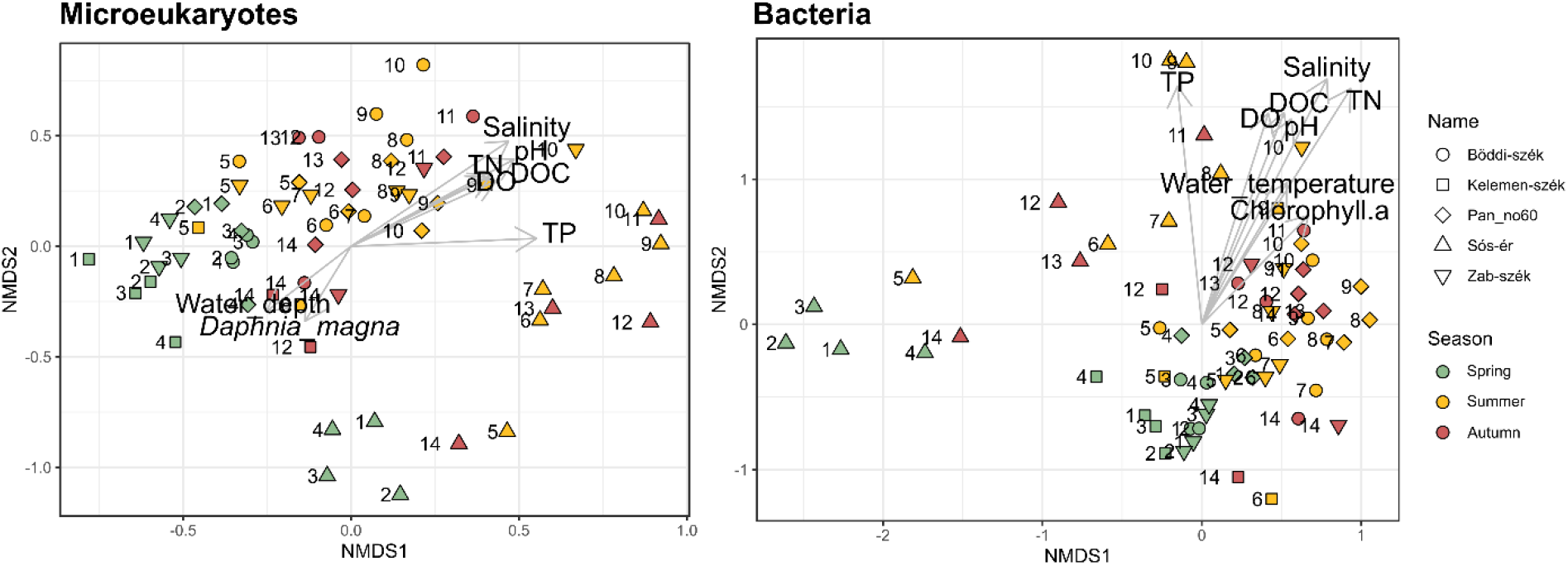
Non-metric multidimensional scaling (NMDS) ordination with the significantly fitted environmental parameters of the microeukaryotic and bacterial planktonic communities of the five soda pans

Meanwhile, the two-way PERMANOVAs testing only the turbid pans showed that for all OTUs or only core OTUs for microeukaryotic communities the season of sampling explained more variance (all OTUs: 20.6%, core: 22.6%) than the identity of the pans, while for bacterial communities the variances explained by the two factors were similar. However, for the non-core communities lake identity explained more variance than season of sampling for both Microeukaryotes and Bacteria. Finally, for both core5 and core4 communities seasonality explained more variation than for the respective non-core communities (Table 1).

**Table 1.**
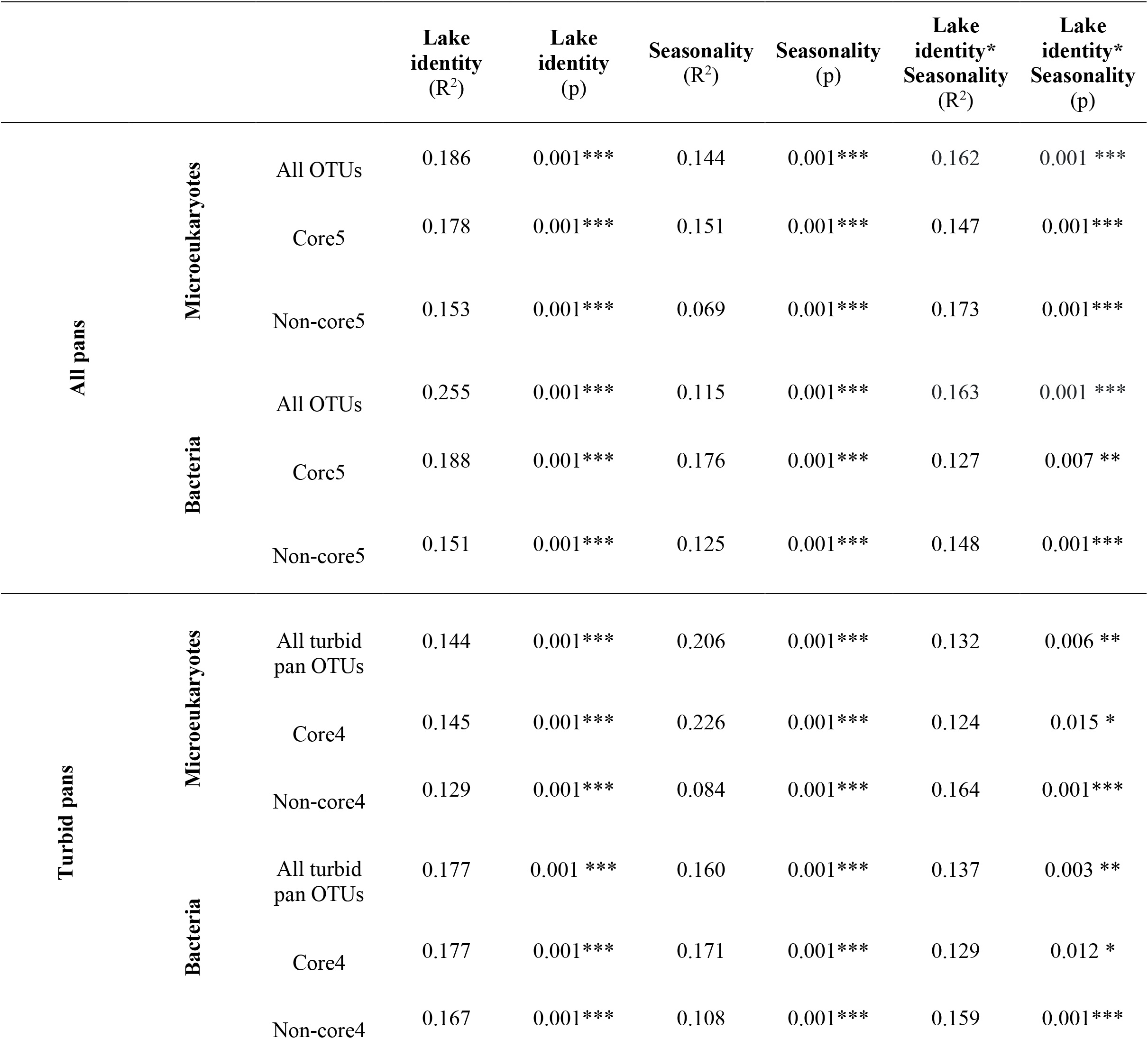
Impact of lake identity and seasonality on the structure of microeukaryotic and bacterial communities (number of * indicates the statistical significance with p, 0 ‘***, 0.001 ‘**’, 0.01 ‘*’, 0.05 ‘.’, 0.1 ‘ ‘, 1)

### Community turnover

Microeukaryotic communities of the turbid pans showed mostly low Bray-Curtis (BC) dissimilarity (< 0.5) between the spring sampling times (April-May) indicating low turnover, while between late spring and early summer BC dissimilarity increased substantially for Pan no. 60, Zab-szék and Kelemen-szék and gradually for Böddi-szék, indicating increasing turnover and a major shift in community structure between the two seasons. Although Zab-szék and Pan no. 60 had low BC dissimilarity in June-July showing a stable period in the first half of summer, starting from mid July to the end of the study period BC dissimilarity remained high (> 0.5) in all pans suggesting high turnover. In the case of the temporary desiccated Zab-szék and Kelemen-szék, the BC dissimilarity was even higher before and after each desiccations, indicating that the microeukaryotic community composition after the refillment was drastically different from the communities before the drought. The BC dissimilarity trend in Sós-ér was different from the turbid pans as it had high values throughout the study period indicating constant high turnover (Figure 5a).

**Figure 5.**
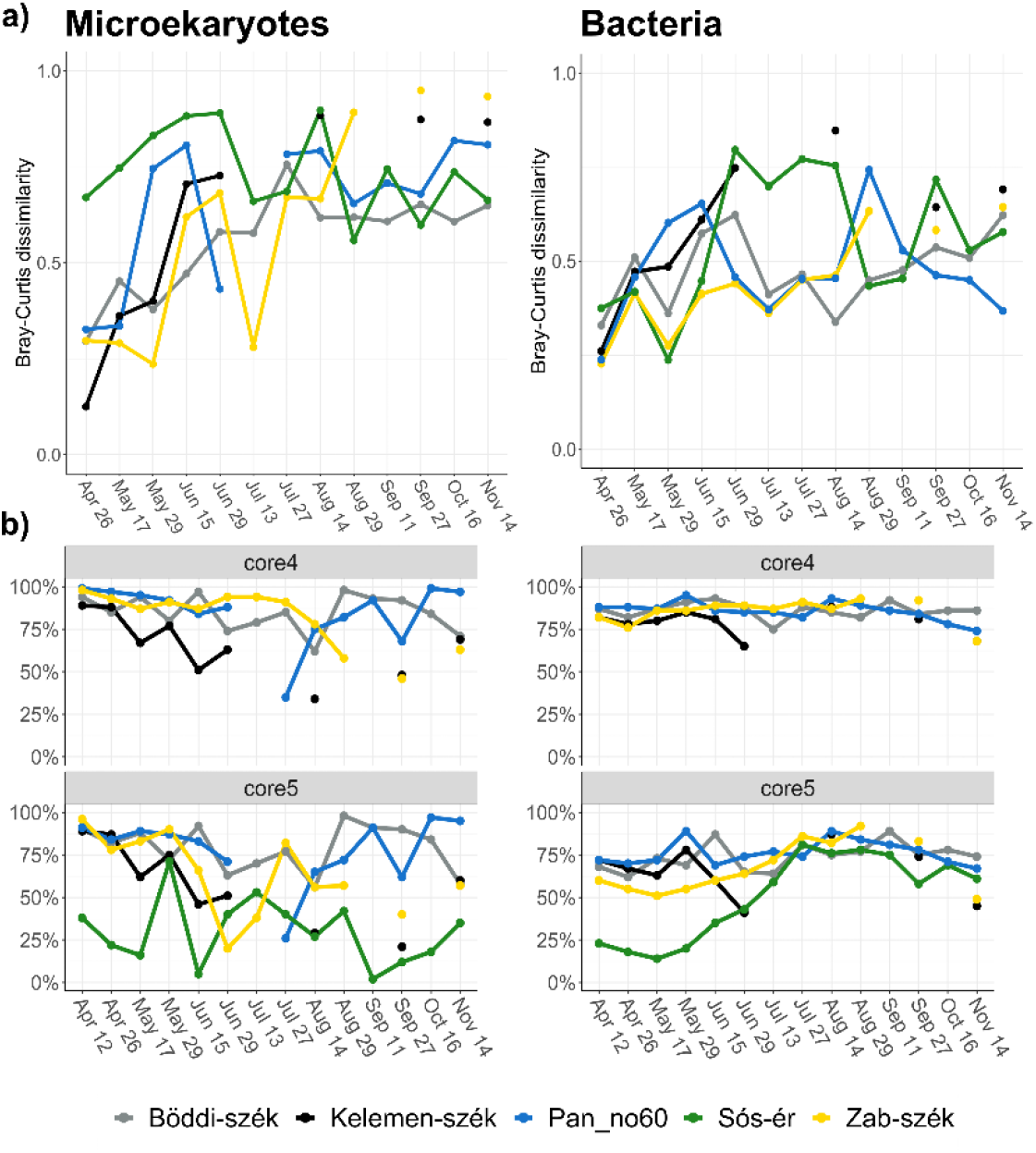
a) Bray-Curtis dissimilarity index between subsequent samplings as proxy for turnover (The X axis represents every second date of the sampling), b) Relative abundance of the core4 and core5 OTUs in the studied pans during the study period

The BC dissimilarity between consecutive samplings was lower for bOTUs (mean 0.5) than for eOTUs (mean 0.6) but the lowest bacteria BC dissimilarities were also measured during spring (April-May) similar to the microeukaryotes. Turnover increase between spring and summer occurred only for Sós-ér and Kelemen-szék, the turbid pan with the most desiccation events. The other turbid pans maintained relatively low turnover throughout the summer, BC dissimilarity increased only in Zab-szék before it dried up. Opposite to the microeukaryotic communities bacterial BC dissimilarity did not increase as a consequence of desiccation events in late summer and in autumn. The only desiccation with turnover increase was the first drought of Kelemen-szék. Opposite to most turbid pans, the bacterial turnover of Sós-ér was high from early summer to mid August. In autumn, each of the three pans that did not dry up followed different trajectories either with decreasing (Pan no. 60), increasing (Böddi-szék) or varying trends of dissimilarity (Sós-ér) (Figure 5a).

### Core microbial community

From the almost 9000 identified OTUs only 97 eOTUs and 191 bOTUs were detected in all five pans, although these core5 OTUs represented 61.8% and 66.8% of the 18S and 16S rRNA gene reads, respectively. These core community OTUs had the highest relative abundance within the microbial communities of each site. The microbial community composition of Sós-ér, the only brown soda pan in this study substantially differed from the other four turbid pans (Böddi-szék, Kelemen-szék, Pan no. 60 and Zab-szék), therefore, we also investigated the OTUs shared by only the four turbid pans (i.e., core4). Core4 consisted of 5-10 times more OTUs than core5 (952 eOTUs and 988 bOTUs), which represented 80.3% and 84.4% of the respective reads (Figure S2).

Sós-ér had the lowest contribution of core5 reads to both its microeukaryotic and bacterial community with 30% and 51%, respectively, followed by the two turbid pans that undergone desiccation (Kelemen- and Zab-szék with 58% and 64% of core5 eOTU reads and 65% and 67% of core5 bOTU reads, respectively). Finally, the highest contribution of core5 was in the permanent turbid lakes (Böddi-szék and Pan no. 60) with a contribution of 79% for eOTU and 75% for bOTU reads. Core4 contribution showed a similar pattern with the lowest ratio in Kelemen-szék (65% eOTU and 79% bOTU reads), the pan that dried out the most times and for the longest periods. Meanwhile, in the other three turbid pans core4 bOTU reads had 86% and core4 eOTU reads had a contribution of 82% in Zab-szék, and 85% in Böddi-szék and Pan no. 60. When it comes to trends over time in Sós-ér, the contribution of core5 eOTUs to the communities was low over the entire study period, while core5 bOTUs had a low contribution only during spring and early summer. In the turbid pans, both core5 and core4 eOTU reads went markedly down after drought events and in the middle of summer in Pan no. 60, while core5 and core4 bOTUs showed a relatively stable contribution over time (Figure 5b).

### Microbial interactions

All networks had more positive correlations than negative irrespective of the pan or type of network (Table S4). Contrastingly to the networks of the non-drying turbid pans (i.e., Böddi-szék and Pan no. 60), the time-shifted networks of the brown Sós-ér and the two turbid pans that underwent several drying-rewetting cycles (i.e., Kelemen-szék and Zab-szék) had less edges and nodes than their synchronous networks. The networks of Sós-ér further distinguished from all turbid networks as they had the lowest numbers of edges, nodes, and neighbours, and together with the time-shifted network of Zab-szék had the lowest density (Table S4). The network topology also showed clear differences between the pans (Figure 6). Both time-shifted and synchronous networks of Böddi-szék and Pan no. 60 had two distinct hubs densely connected with mostly SSCC edges (i.e., global correlations). Meanwhile, the nodes of synchronous networks of Kelemen-szék and Zab-szék were densely connected with mostly LS edges (i.e., local correlations). The topology of the time-shifted association network of Zab-szék and both networks of Sós-ér were the most fragmented, for Sós-ér even an additional hub corresponding to the *Nodularia* bloom period was distinguishable (Figure 6).

**Figure 6.**
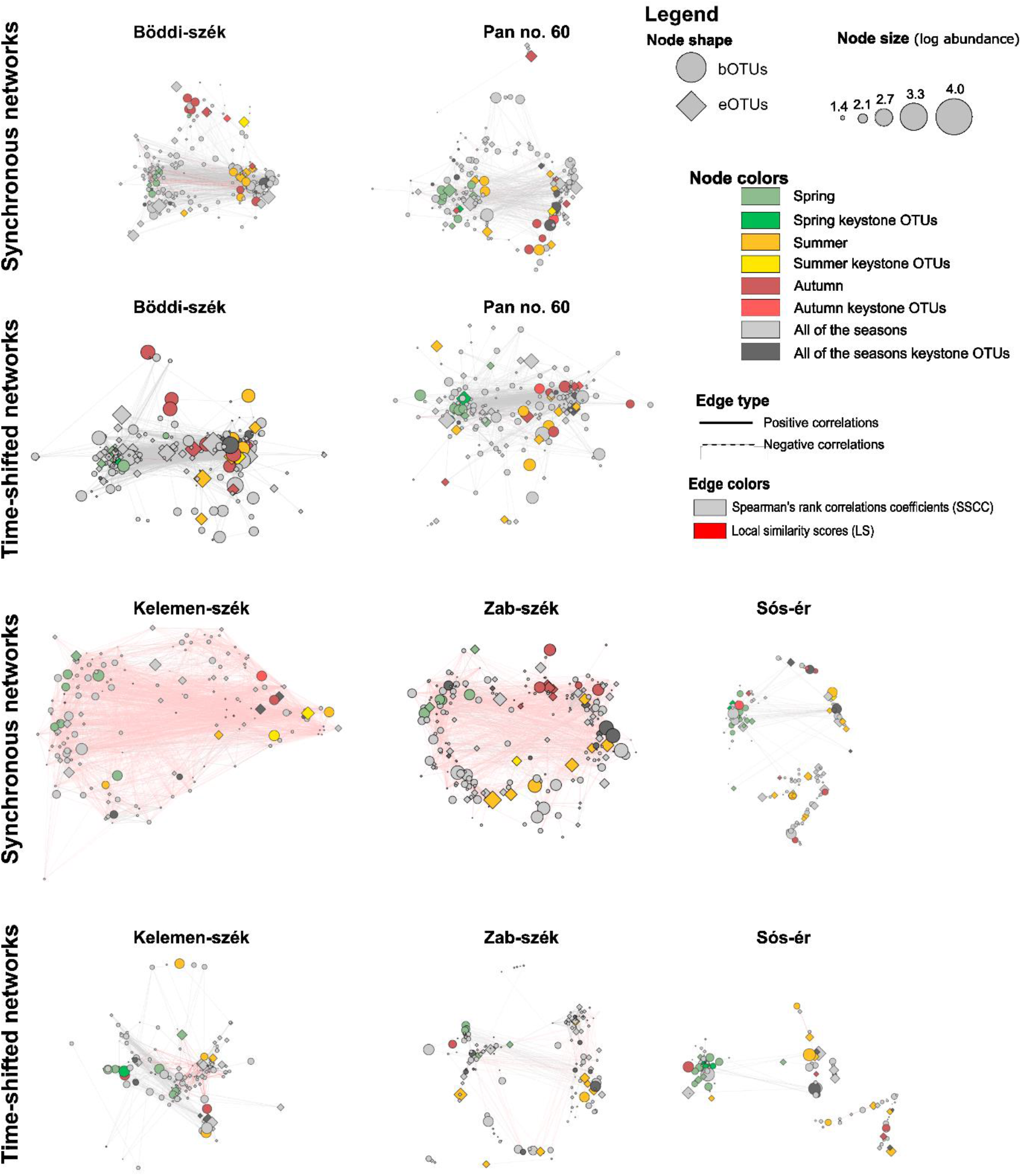
Synchronous and time-shifted networks of bacterial and microeukaryotic OTUs of the soda pans. The colouring of the network nodes was based on the difference of the mean abundance of the corresponding OTUs in spring, summer or, autumn: green = deviation of the mean abundance of the sampling period < mean abundance in spring nodes yellow = deviation of the mean abundance of the sampling period < mean abundance in summer nodes red = deviation of the mean abundance of the sampling period < mean abundance in autumn nodes grey = deviation of the mean abundance of the sampling period > mean abundance of spring, summer and autumn nodes. The colouring of the key OTUs are brighter than the other colours.

Although microeukaryotic and bacterial OTUs were both identified as keystone OTUs in all networks, the majority of keystone OTUs were bacteria and there were more bacterial keystone OTUs in the turbid pans than in Sós-ér (Figure 7).

**Figure 7.**
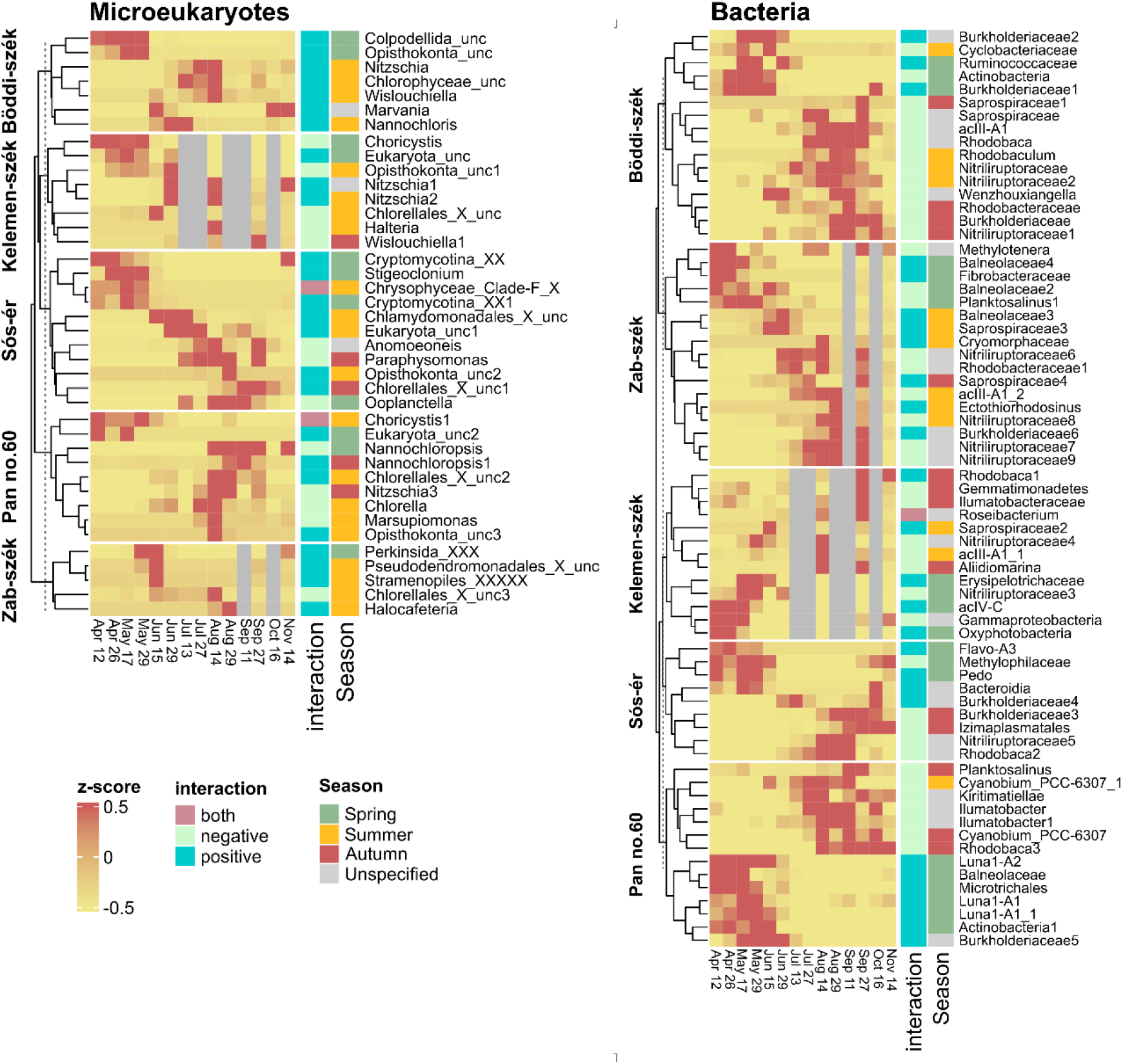
Taxonomic affiliation and distribution of the top keystone OTUs of the networks based on their z-score

Only one-third (36%) of eukaryotic keystones was core5, while for the turbid lakes 58% of s were core4. Among the keystone bOTUs the majority was core5 (79%) and almost all were core4 (91%) in the turbid lakes. Many keystone OTUs were assigned to phyla with high relative abundance such as Chlorophyta, Ochrophyta, and Fungi for keystone eOTUs and Actinobacteria for bOTUs. However, Cyanobacteria were underrepresented among keystones with even very abundant taxa such as *Nodularia*_PCC-9350 and *Synechococcus*_MBIC10613 not identified as a keystone. Furthermore, the highly abundant Erysipelotrichaceae OTU in Sós-ér, was also not a keystone. Meanwhile, among the less abundant (<0.1% of reads) keystones there were two eOTUs assigned to the parasitic Cryptomycotina order.

There were various differences among positive and negative keystone OTUs (Figure 7). First, positive keystones were more common among those with highest abundance in spring (74% of spring keystones), in summer the ratio of positives and negatives was similar, while among those abundant in autumn negatives were more common (80%). For bOTUs, taxonomic differences between positives and negatives were also noted. On the phylum level Bacteroidetes (10) was the most common among positive keystones followed by Proteobacteria (7) and then Actinobacteria (6), while for the negative OTUs Actinobacteria was the most common (17), followed by Proteobacteria (13) and then Bacteroidetes (6). Differences were obvious also at lower taxonomic levels as negative Actinobacterial keystones were present in high abundance throughout the study period and primarily belonged to the Nitriliruptoraceae (10) and acIII-A1 (3) lineages, while the positives were only abundant in spring and belonged to Luna1-A (3), and acIV-C (1). Similarly, the most common family for negative Proteobacterial keystones was summer and autumn abundant Rhodobacteraceae, while for positives it was Burkholderiaceae (5) in all three seasons.

No substantial taxonomic difference was notable between keystone eOTUs of the synchronous and time-shifted networks. However, for bacterial keystones Actinobacteria were more common in the synchronous networks than in the time-shifted ones (16 vs. 8), while Bacteroidetes keystones were more common in time-shifted networks (10 vs. 6).

## Discussion

Planktonic microbial communities of five shallow soda lakes exposed to identical climatic and meteorological conditions were evaluated by a synchronous time series analysis. The results demonstrated that the pans shared a core microbial community and similar seasonal dynamics both in respect of community composition and interactions. However, the extent of shared microbiome and trends were not uniform across the lakes. Substantial differences were identified based on habitat subtype (i.e., brown or turbid). Common seasonal succession trajectories were also disrupted by local stochastic events (e.g., desiccation). Such events prompted stronger response from microeukaryotic than from bacterial communities.

Although, both the brown and turbid pans investigated in this study represent characteristic soda pan habitats (i.e., high pH, specific ion composition, shallowness) ^19^, the brown Sós-ér is also characterised by extremely high CDOM (coloured dissolved organic matter) and DOC concentrations, and abundant shoreline vegetation ^10^. Furthermore, it has been previously demonstrated that Sós-ér harbours markedly different microbial communities than the turbid Zab-szék ^47,69^. During our study period, Sós-ér also had significantly deeper waters, and higher TN concentrations as well as significantly different microeukaryotic and bacterial communities than the turbid pans. These differences were reflected in a substantially lower number of shared OTUs among all five pans (core5) than among the turbid pans (core4). The contribution of core5 eOTUs to Sós-ér community was low throughout the study period and various unique microeukaryotic taxa were detected with high relative abundance only in this pan, indicating that different soda pan subtypes exert strong selective forces on microeukaryotes. Meanwhile, from mid summer the contribution of core5 bOTUs to Sós-ér community was similarly high as in the turbid pans, suggesting for bacteria more similar selection processes and no dispersal limitation between the five lakes. Sós-ér also had various strikingly different intrinsic characteristics such as fragmented, low density and connectivity networks as well as high turnover of eOTUs that indicate highly dynamic communities. Interestingly, the bOTUs of Sós-ér had high turnover only during summer and autumn, while in spring bacterial turnover was low suggesting, together with the low core5 bOTU contribution in this period, a distinctive but stable spring bacterial community. This community was characterised by high abundance of *Erysipelotrichaceae* UCG-004 bOTU, a taxon that was previously reported from soda lakes of this region ^47^ but has been otherwise described primarily from the digestive systems of mammals and insects ^70–72^.

Seasonality significantly affected the microeukaryotic and bacterial communities of all five pans and all studied seasons had distinctive communities. Furthermore, despite the differences of Sós-ér, various common attributes of seasonal microbial succession could be identified. First, a circular trajectory of seasonality indicated by the resemblance of early spring and late autumn communities was detected for all pans, which was driven by low water temperature and deeper waters and consequently low pH and low concentration of solutes. The seasonal variation explained by the core vs. non-core communities also showed similarities irrespective of including Sós-ér in the analyses (i.e., core5 or core4). More specifically, the higher seasonal variance explained by the core than by the respective non-core communities supported our hypothesis in respect of seasonal adaptation happening primarily through species recruitment from the core community. Although, not surprisingly, the variation explained by the identity of lakes was higher for the five lakes than for turbid pans only.

Further similarities of the seasonality of the five pans were the analogous attributes of spring. Spring was the season that differentiated the most strongly from the others (i.e., summer and autumn) both according to community structure and interactions. In all five lakes, spring was characterised by positive keystones. Positive synchronous associations can reflect mutualistic and facilitative interactions but also parasitism, predation or similar niche preference. However, the direction of species correlations is not always obvious, for example similar niche preference can manifest both as positive synchronous correlation due to coexistence or negative due to competitive exclusion. Similarly, predation can display as positive synchronous associations when a prey attracts its predators to a habitat patch or as negative (both synchronous and time-shifted) when predators eliminate their preys ^73,74^. Various positive keystone eOTUs abundant in spring were assigned to predatory flagellates such as Colpodellida ^75^ or intracellular parasitic taxa such as Cryptomycotina ^76^ and Perkinsozoa ^77^ which are common in aquatic habitats ^78^. These, together with the significant effect of *Daphnia magna* on microeukaryotic community structure suggest an important role of top-down controls in spring. Meanwhile, the spring abundant positive actinobacterial keystones belonged to lineages (i.e., Luna1-A and acIV-C) that are characterised by very small cell-sizes (< 0.1 µm^3, 79^ and have been previously suggested to be grazing resistant ^80,81^ which might explain their coexistence with predatory eukaryotic taxa. Overall, the seasonal dynamics of actinobacterial keystones were similar to seasonal dynamics described for this phylum in various other limnic systems ^56,82^.

Except for the eOTUs of Sós-ér, spring was also characterised by low species turnover, which could be the result of relative stability provided by deeper waters. The spring samples had high abundance of a single *Choricystis* eOTU, a widely distributed freshwater picoeukaryotic green algal genus ^83^. Our results are in agreement with previous studies showing that in soda pans of the same region picoeukaryotic green algae are the most abundant members of the phytoplankton and have a specific seasonal trend with the highest abundances in winter-spring and the lowest in summer ^40,45,47,84,85^. The dominance of positive keystone taxa and the transitivity of positive correlations combined with low turnover of both eOTUs and bOTUs in the turbid lakes suggests that spring was a period when community assembly was primarily ruled by trophic interactions between primary producer eukaryotic picoalgae (e.g., *Choricystis*), their parasites (e.g., Cryptomycotina), heterotrophic bacteria (e.g., Luna1 lineages, Burkholderiaceae, Balneolaceae) consuming algal exudates and debris, and flagellates (e.g., Colpodellida) predating on bacteria.

After the relatively stable and synchronous period in spring, the variation between microbial communities increased even for the turbid pans. This was driven by the higher and more variable concentrations of dissolved substances resulting from the lower water levels which is in agreement with previous studies showing that environmental fluctuations induced by shrinking ecosystem size modulate community assembly processes and reduce stability ^86^. In the case of Kelemen-szék and Zab-szék the decreasing water levels resulted in various desiccation events followed by refillments. Drying-rewetting cycles exert severe stress to microorganisms due to drastic changes in salt and nutrient content ^87–89^. While desiccation is common in soda pans of this region, not every pan dries out every year and it is not always the same pans that dry out ^10,47^ making desiccation not part of the regular seasonality but rather a local stochastic stressor.

Intense environmental fluctuation combined with increased growth rates due to summer warming combined with decreasing habitat size (i.e., shrinking water levels) was expected to stimulate microbial turnover ^86,90^. Accordingly, the turnover of microeukaryotic communities in the turbid pans increased relatively uniformly. Microeukaryotic turnover also increased substantially as a consequence of each drying-rewetting cycle suggesting limited resilience of the microeukaryotic communities to such stochastic stress. Simultaneously, the contribution of non-core4 eOTUs increased following drought events, supporting our hypothesis about non-core OTUs being more important in the response to stochastic events.

Meanwhile, the turnover of the bacterial communities of the turbid pans remained relatively similar through the study irrespective of desiccation. Bacterial turnover increases were only detected when one of the compared samplings happened at extremely low water levels (≤ 2 cm) suggesting that extremely shallow conditions implied strong selective force on bacterial communities but bacteria were otherwise more resistant to the desiccation stress and the overall more extreme conditions of summer-autumn than microeukaryotes. Bacterial keystone of this period also suggests special adaptations to extreme conditions. For example, it has been shown for close relatives of the summer-autumn abundant Nitriliruptoraceae keystones that they can scavenge organic nitrogen even from strong nitrile bonds ^91^ allowing them versatility to overcome nitrogen-limitation (Neuenschwander et al., 2017; A. Szabó et al., 2020). The high number of negative bacterial keystones in summer-autumn also suggests that bacterial groups with different optima were dynamically outcompeting each other under the quickly changing conditions. The contribution of core4 bOTUs was also very high at every sampling time and site suggesting that the core bacterial community of turbid pans not only contributed to the adaptation to seasonal changes but is also highly resistant to extreme conditions. As it has been shown that drying-rewetting cycles have strong filtering effects on bacterial communities and dispersal is required for full recovery ^88,93,94^, the uninterrupted high contribution of core4 bOTUs irrespective of drought events suggests no dispersal limitation for bacteria between the studied pans. Recently, it has been suggested that waterbirds play an important role in dispersing both prokaryotes and microeukaryotes between soda pans (i.e., endozoochory) ^48^, which implies that dispersal intensity would vary with bird visitation frequency. Although, our study covered periods with different bird abundances ^44^, the absence of indications of bacterial dispersal limitation suggests that the bacterial core microbiome is highly dispersed between the pans by non-endozoochory dispersal such as wind or precipitation ^95^.

In summary, these results together with the higher contribution of core bOTUs to keystone taxa than that of core eOTUs, indicate more synchronised community trends for bacteria than for microeukaryotes. In general, microeukaryotic communities were more sensitive to stochastic events, probably both due to lesser physiological resistance and dispersal limitation. All in all this supports our hypothesis regarding the lesser impact of stochastic events on bacterial compared to microeukaryotic communities. Our results are also in agreement with studies suggesting that dispersal limitation and stochasticity are more important in shaping microeukaryotic communities, while selection processes are more prominent in bacterial community assembly and with studies that identified increased dispersal limitation during dry periods for microeukaryotic communities (Beisner et al., 2006; Chen et al., 2019; Logares et al., 2018; Mikhailov et al., 2022; Y. Wang et al., 2015). We also demonstrated that while both the microeukaryotic and bacterial communities were driven by similar physical and chemical factors, zooplankton had significant effect only on eOTUs indicating top-down control, while chlorophyll only affected bOTUs suggesting greater importance of bottom-up control for the primarily heterotrophic bacteria.

The networks generated in this study allowed for joint analyses of the interactions of microeukaryotes and bacteria. All networks had more positive correlations than negatives suggesting predominance of positive interactions in the communities. Previous studies demonstrated that positive associations are more common in ecosystems characterised by high abiotic stress due to higher number of mutualistic interactions permitting species to exist in harsher environments than would otherwise be possible ^98,99^. However, the dependence on mutualism makes such communities sensitive to perturbations and reduces network stability particularly in case of low modularity networks ^99^. Appart of the dominance of positive correlations, there were clear differences in network topology and properties between the brown Sós-ér, the two drying turbid pans (Kelemen-szék and Zab-szék) and the turbid pans that did not experience drought (Böddi-szék and Pan no.60). The synchronous networks of the non-drying turbid pans had two distinct hubs corresponding to the spring and summer-autumn period, while the networks of drying pans had less clear modularity and networks of Sós-ér had three hubs with one corresponding to the late summer period characterised by *Nodularia* bloom. The hubs of non-drying turbid pans were connected with mainly negative SSCC associations suggesting more stability across the entire study period, which was also supported by higher number of correlations and nodes in their time-shifted than in their synchronous networks. Meanwhile, synchronous networks of the drying pans were dominated by LS edges indicating the lack of community dynamics overarching the entire study period, which was further corroborated by lower number of time-shifted than synchronous associations. In summary, the interaction networks of the soda pans reflected low community stability in these high stress habitats that was further exacerbated by the stochastic stress events of drying-rewetting cycles or cyanobacterial blooms.

## Conclusions

By expanding community composition explorations with network analyses and the identification of keystone taxa, we were able to demonstrate unique attributes of the seasonality of extreme aquatic habitats. Namely, we showed that the studied soda pans despite drastic environmental changes and subsequent community shifts are primarily populated by a common core microbiome and as a consequence of the identical climatic and meteorological conditions share certain characteristics of their seasonal trends. However, the extent of the shared microbiome was curtailed among pans of different habitat subtype (i.e., brown or turbid pan) and common seasonal trends were modified by local stochastic environmental events such as desiccation and refillment. In general, the stable environmental conditions of spring fostered stable microbial communities ruled by trophic interactions, while the dynamically changing conditions of summer and early autumn combined with stochastic events exerted strong selective forces that instigated different response mechanisms for microeukaryotic and bacterial communities supporting our hypothesis about microeukaryotic communities being more sensitive to stochastic effects than bacteria. Adaptation to uniform seasonal changes was through species recruitment from the core community, while for microeukaryotes non-core members of the microbiome were involved in the response to stochastic events suggesting that dispersal limitation might have modulated their recovery. Meanwhile, bacterial communities were resistant to stochastic events and adapted to extreme conditions via species sorting from the core community and competitive exclusion. Overall, our results indicate the importance of considering microeukaryotic and bacterial communities separately when it comes to stochastic stressors, particularly as events such as desiccation and refillment become more frequent in shallow aquatic ecosystems due to climate change.

## Acknowledgements

The authors thank Nóra Tugyi, Balázs Németh, Tímea Szabó (Balaton Limnological Research Institute, Hungary) and Emil Boros (Institute of Aquatic Ecology, Centre for Ecological Research, Hungary) for their help in fieldwork and measurements of chlorophyll a; Christoffer Bergvall (Uppsala University, Sweden) for laboratory assistance in the measurement of TP, TN and DOC concentrations; Tamás Sápi (Kiskunság National Park) for help in fieldwork; Zsófia Horváth (Institute of Aquatic Ecology, Centre for Ecological Research, Hungary) for help and training in microscopic identification of zooplankton; and the Hungarian Meteorological Service for providing the meteorological data.

## Data Availability Statement

Raw sequence reads can be accessed at the NCBI SRA through BioProject ID PRJNA272672 and Biosample IDs SAMN32532920-SAMN32532982.

## Contributions

TF, ASz, AJSz conceived the study; ZsM, BCs, TF, ASz collected the samples; ZsM, BCs, conducted laboratory work under supervision of AJSz; ZsM, ASz, AJSz, FJ and AH performed all data analysis; ZsM, ASz, AJSz contributed to data interpretation; ZsM wrote the manuscript with the guidance and revision of AJSz and ASz, and all authors contributed the manuscript final version.

