## Supplementary material for "Contrasting Response of Microeukaryotic and Bacterial Communities to the Interplay of Seasonality and Stochastic Events in Shallow Soda Lakes"

This file includes the following text, tables and figures:

Text S1-S3

Table S2-S4

Figure S1-S2

**Text S1** 18S and 16S rRNA gene amplicon sequencing

Eukaryotic primers 574*F (CGGTAAYTCCAGCTCYAV) and 1132R (CCGTCAATTHCTTYAART) [1] and prokaryotic primers 341F (CCTACGGGNGGCWGCAG) [2], 805NR (GACTACHVGGGTATCTAATCC) [3] were used for the polymerase chain reactions. To decrease the stochastic effect of the reaction, all PCR amplification was performed in duplicates in 20 µL, which contained 4 µL of 5x Q5 reaction buffer, 2 µL of dNTP (2 mM), 0.2 µL of Q5 High Fidelity DNA polymerase (2 U/µL) (New England Biolabs), 0.5 µL of each primer (10 µM), 11.8 µL of nuclease free water and 1 µL of template DNA. The following thermal cycle conditions were used for 18S rRNA gene amplification: initial denaturation at 98 °C for 1 min 10 sec, followed by 20 cycles (annealing at 51 °C for 30 sec, extension at 72 °C for 30 sec) and a final elongation step at 72 °C for 2 min. The following thermal cycle conditions were used for 16S rRNA gene amplification: initial denaturation at 98 °C for 40 sec, followed by 20 cycles (annealing at 48 °C for 30 sec, extension at 72 °C for 30 sec) and a final elongation step at 72 °C for 2 min. Amplicons were pooled before purification with magnetic beads (Agencourt AMPure XP PCR Purification, 2013). To prepare libraries for Illumina sequencing, primers were prolonged by illumina handles and index primers. The second PCR reaction contained 4 µL of 5x Q5 reaction buffer, 2 µL of dNTP (2 mM), 0.2 µL of Q5 High Fidelity DNA polymerase, 1 µL of each index primer (5 µM), 9.8 µL of nuclease free water and 1 µL of template from the first PCR. The following thermal cycle was used for both eukaryote and prokaryote specific reaction: initial denaturation 98 °C for 40 sec, followed by 15 cycles of denaturation 98 °C for 10 sec, annealing at 66 °C for 30 sec, extension at 72 °C for 30 sec/kb and the final extension at 72°C for 2 min. Amplicons were purified again with magnetic beads (Agencourt AMPure) as described previously. Quantification of the libraries were carried out using a PicoGreen assay (Quant-iT PicoGreen dsDNA Assay Kit, Invitrogen). Sequencing was performed at the SciLifeLab (Uppsala, Sweden) on an Illumina MiSeq platform (Illumina Inc, San Diego, CA, USA).

**Text S2 Bioinformatic analysis of the sequencing data**

Bioinformatic analysis of the sequence reads were carried out with mothur v1.41.1[4] using the MiSeq SOP (http://www.mothur.org/wiki/MiSeq_SOP downloaded at 9th July 2018). The deltaq parameter of the ’make.contigs’ command was adjusted to 10 for additional quality filtering to eliminate sequencing errors. Primers were removed from the start and the end of the sequences and singletons were also removed from the dataset according to [5]. For the alignment of sequence reads the ARB-SILVA SSU Ref NR 132 reference database [6] was used. Denoising was performed using mothur's pre.cluster command using the default algorithm (Huse et al., 2010) and applying the suggested 4 bp difference cutoff. Chimeras were identified and removed using the mothur implemented version of VSEARCH. Operational taxonomic units (OTUs) were assigned at 99% similarity threshold levels with the OptiClust algorithm [7]. Taxonomic assignment of the 18S rRNA gene OTUs was carried out using the PR$^2$ v4.10 reference database [8] with a minimum bootstrap confidence score of 80 and applying 1000 iterations. For the 16S rRNA gene amplicon set the TaxAss software [9] was used for taxonomic classification applying default parameters and using the FreshTrain (2018 April 30 release) and ARB-SILVA SSU Ref NR 132 databases as reference. Bacterial OTUs assigned to non-primer specific taxonomic groups (e.g. Archaea, chloroplasts, mitochondria, unknown) and microeukaryotic OTUs assigned to taxa Metazoa, Streptophyta, Basidiomycota and Ascomycota (due to the prefiltration through a 40 µm pore sized mesh of the water samples) were removed from the dataset. The 7th sampling time of Pan no. 60 was discarded from the 18S rRNA gene amplicon dataset due to low number of high-quality sequences. For statistical analyses, reads were subsampled to the read number of the sample having the lowest sequence count (62 samples in the 18S rRNA amplicon set, n = 2407 and 63 samples in the 16S rRNA amplicon set, n = 3188).

**Text S3 Measured environmental parameters**

Water temperature increased from 13.7 °C to 27.8 °C during spring (mean: 20.0 °C) and decreased from 24.8 °C to 7.1 °C during autumn (mean: 16.3 °C), while in summer varied between 19.6 and 30.9 °C (mean: 25.2 °C). Salinity values varied between the subsaline (min. 0.9 g/L) and mesosaline (max. 27.8 g/L) categories with the majority of samples (54 out of 63) being hyposaline (3-20 g/L) [10]. The values of DOC, TN and TP varied between 10 and 3341 mg/L, 1.5 and 25.7 mg/L and 0.5 and 25 mg/L, respectively. The pH value remained alkaline throughout the study period in all pans with an average of 9.5 pH and varying from 8.5 to 10.0. The pans were aerobic at each sampling time ($O_2$ saturation >79%) and often over-saturated ($O_2$ saturation >100%). Chlorophyll a concentration ranged between 1.8 and 696.7 µg/L, with an average of 204.1 µg/L with both the lowest and highest chlorophyll a values measured in Pan no. 60 (Figure S1).

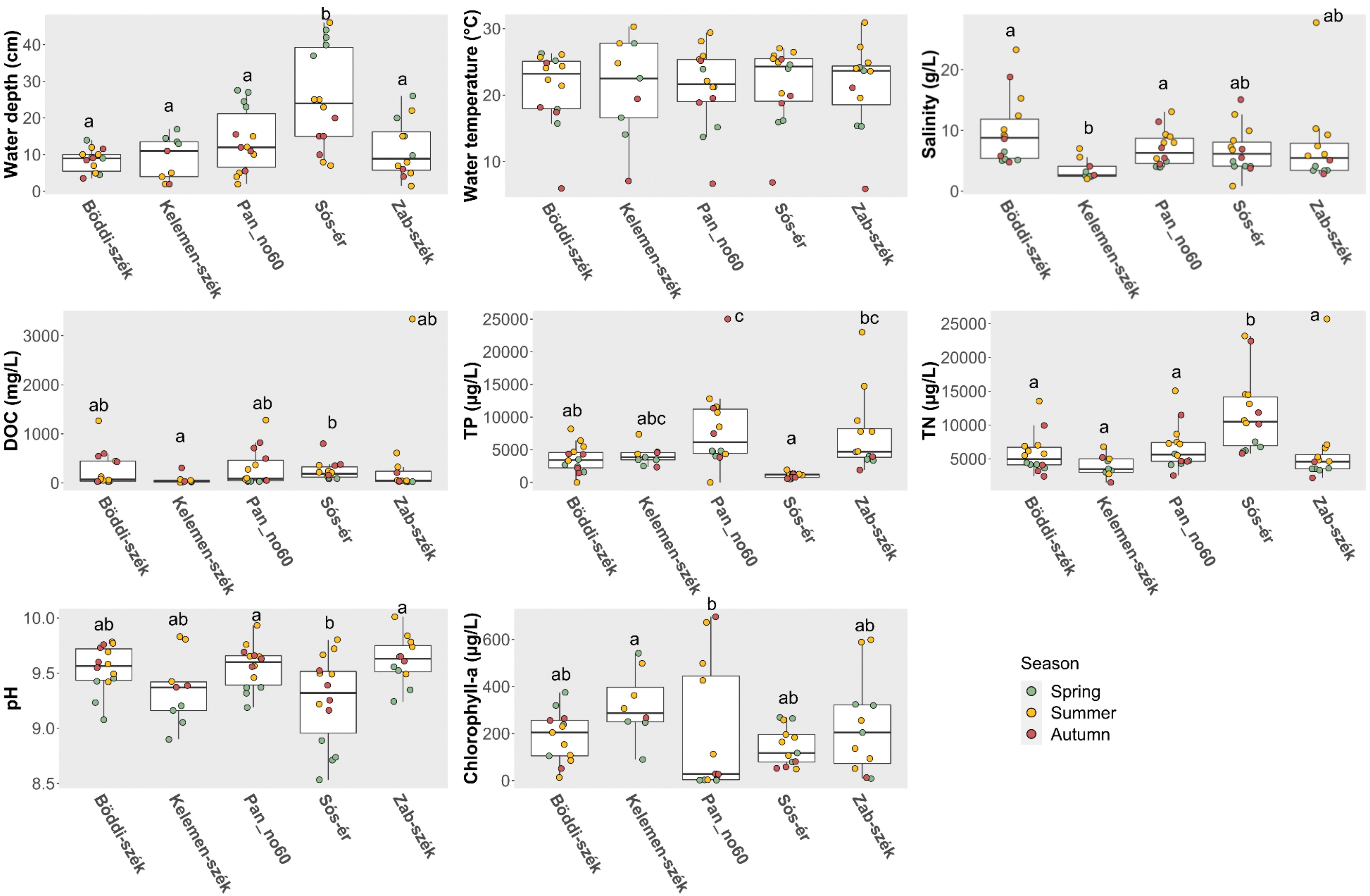


**Figure S1** Environmental parameters of the soda pans. Different letters within soda pans indicate statistically significant differences at a significant level of $p < 0.05$.

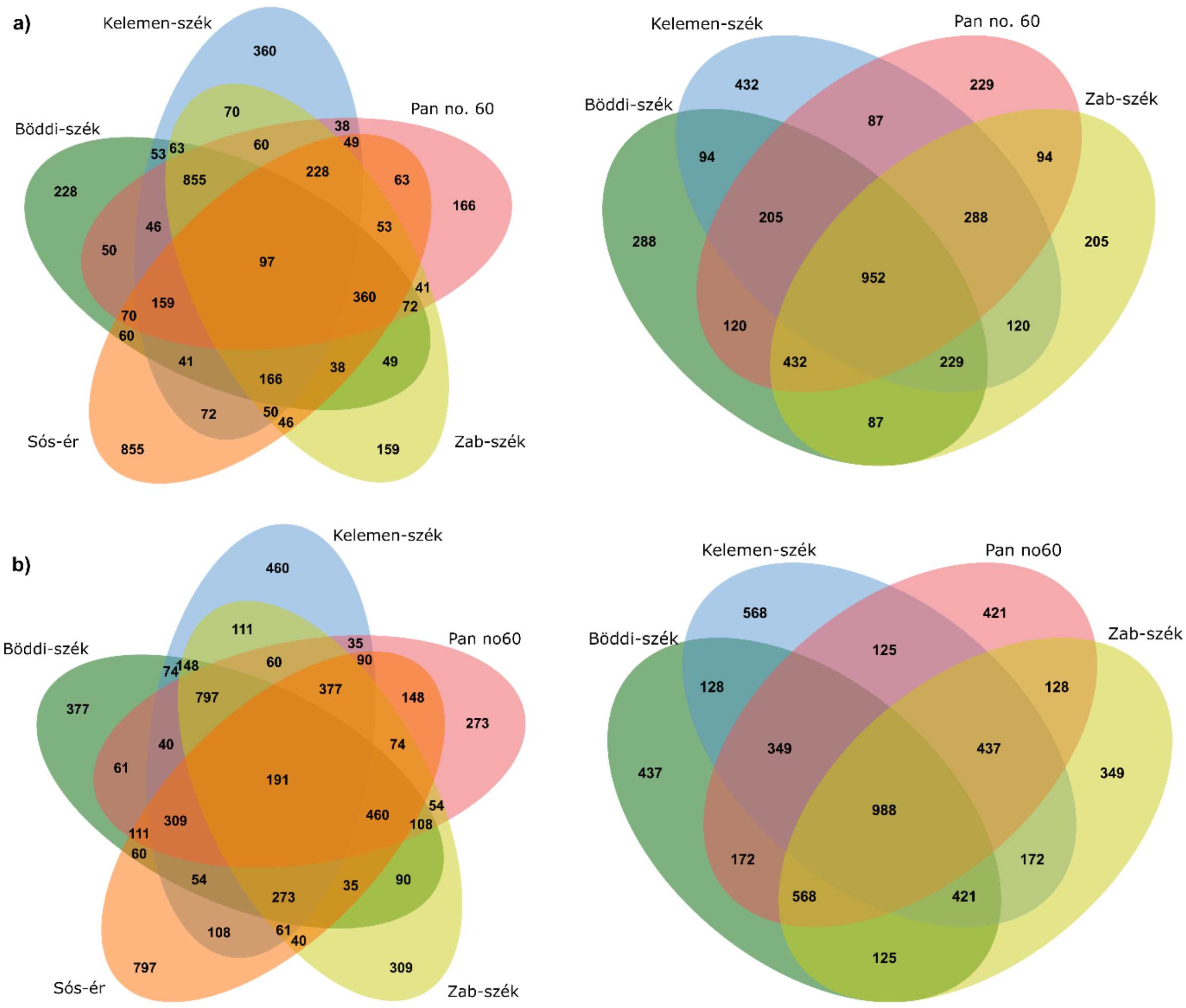


**Figure S2** Venn diagrams showing the numbers of shared and unique OTUs between all the five lakes and the four turbid type of soda pans: a) Microeukaryotic OTUs, b) Bacterial OTUs (Venn diagrams were generated using jvenn[11], to visualize the core microbial community of the pans)

**Table S3** Differences of planktonic microbial communities from the five soda pans comparing the three studied seasons. (PERMANOVA test results; number of * indicates the statistical significance with p. 0 ’***. 0.001 ’**’. 0.01 ’*’. 0.05 ’.’. 0.1 ’ ’. 1)

| | Spring/Summer ($R^2$) | Spring/Summer (p) | Spring/Autumn ($R^2$) | Spring/Autumn (p) | Summer/Autumn ($R^2$) | Summer/Autumn (p) |
|---|---|---|---|---|---|---|
| **Micoreukaryotes** | 0.129 | 0.001*** | 0.185 | 0.001*** | 0.043 | 0.014* |
| **Bacteria** | 0.102 | 0.001*** | 0.124 | 0.001*** | 0.039 | 0.045* |

**Table S4** General network properties generated from the Network Analyzer of Cytoscape v3.8.2. a) Synchronous b) Time-shifted. (* Number of edges = Number of negative correlations + Number of positive correlations)

| **a)** | **Lake** | **Number of nodes** | **Number of edges*** | **Average number of neighbours** | **Density** | **Number of negative correlations** | **Number of positive correlations** |
|---|---|---|---|---|---|---|---|
| | **Böddi-szék** | 199 | 1417 | 14.24 | 0.07 | 506 | 911 |
| | **Kelemen-szék** | 170 | 2672 | 31.44 | 0.19 | 1020 | 1652 |
| | **Pan no. 60** | 176 | 848 | 9.88 | 0.06 | 304 | 544 |
| | **Sós-ér** | 139 | 314 | 5.17 | 0.05 | 53 | 261 |
| | **Zab-szék** | | | | | | |
| **b)** | | | | | | | |
| | **Böddi-szék** | 202 | 2246 | 22.24 | 0.11 | 911 | 1335 |
| | **Kelemen-szék** | 147 | 689 | 9.37 | 0.06 | 207 | 482 |
| | **Pan no. 60** | 182 | 1304 | 14.55 | 0.08 | 508 | 796 |
| | **Sós-ér** | 99 | 153 | 4.03 | 0.07 | 12 | 141 |
| | **Zab-szék** | 156 | 535 | 7.08 | 0.05 | 85 | 450 |
